# YAMB: metagenome binning using nonlinear dimensionality reduction and density-based clustering

**DOI:** 10.1101/521286

**Authors:** Aleksei Korzhenkov

## Abstract

**Summary:** YAMB is a novel metagenome binning tool, which uses tetranucletotide composition and average contig coverage and performs t-SNE dimensionality reduction and sequential DBSCAN data clusterization. YAMB provided with metagenomics assembly and reads may be used for straightforward metagenome binning on a recent personal computer running Linux OS.

**Availability and Implementation:** Source code of YAMB is freely available on GitHub (https://github.com/laxeye/YAMB), implemented in R, Perl and Bash and supported on Linux.

## 1 Introduction

Modern tools for metagenome contig binning are mainly based on nucleotide composition and relative abundance of sequences such as CONCOCT (Alneberg et al, 2014), MetaBAT (Kang et al., 2015), COCACOLA (Lu et al., 2017), GroopM (Imelfort et al., 2014), MaxBin2 (Wu et al., 2015). These tools perform dimensionality scaling of multimeric data (n>100) and sequential clustering. Nowadays many different methods realized as libraries for several programming languages are available for dimensionality scaling including linear techniques, classical multidimensional scaling, factor analysis and wide spectrum of nonlinear techniques. Despite showing good results on synthetic data these methods may be ineffective for real data (van der Maaten, 2009; Cunningham, Ghahramani, 2015; Sorzano et al., 2014). Such drawbacks affect effectiveness of metagenome binning software.

Method of t-distributed stochastic neighbor embedding (t-SNE) is an evolution of SNE method which implements a heavy-tailed distribution in the low-dimensional space to reduce data crowding and facilitates cost function optimization comparing to SNE (Maaten, Hinton, 2008). t-SNE showed good performance in data analysis and visualization in many applications such as machine learning, image and sound recognition and further improved by Barnes-Hut algorithm (van der Maaten, 2014). Data clustering is another crucial step of metagenome binning, so ability of HDBSCAN (Campello et al., 2015) to be coupled with t-SNE for metagenomic data analysis. HDBSCAN (Hierarchical density-based spatial clustering of applications with noise) generalizes and improves existing density-based clustering techniques providing complete clustering hierarchy composed of all possible density-based clusters following the nonparametric model adopted, for an infinite range of density thresholds. This improvement solves the case of hierarchical algorithms based on density estimates e.g. OPTICS, AUTO-HDS, gSkeletonClu which have a single density threshold that leads to impossibility of simultaneous detection of clusters of varied densities, which is also a major shortcoming several density-based non-hierarchical algorithms e.g. DBSCAN, DENCLUE (Campello et al., 2013).

## 2 Methods

YAMB (yet another metagenome binner) is implemented on several programming languages including Bash, Perl and R as command-line non-interactive application. User provides only metagenome assembly in FASTA format and sequencing reads in FASTQ format. YAMB pipeline includes several steps listed below. Contigs with length less than 1000 bp are discarded. Remaining contigs are cut to the fragments with length maximally close to but not less than 10 000 bp (this value may be adjusted) to increase contribution of longer contigs. This approach was introduced by CONCOCT (Alneberg, 2014). Tetranucleotide occurrences are calculated for each sequence fragment using script written in Perl. Reads may be mapped onto contig fragments using any read-mapping software e.g. bowtie2 (by default), bwa, minimap2. Mean contig coverage is calculated using samtools (Li et. al., 2009) and awk. Data were imported in R environment and threated as matrix where rows corresponded to contig fragments and columns to tetranucleotide occurrences and fragment mean coverage. Dimensionality scaling with t-SNE is performed on normalized data with three distinct perplexity values proportional to total fragment number. Subsequent clustering with HDBSCAN was performed. Binning results are saved as comma-separated values and visualized as two dimensional plots and saved in PNG format using ggplot2 library. Sequences belonging to distinct clusters are extracted and metagenomics bins in FASTA format are written. Completeness and contamination assessment of resulting metagenomic bins by appropriate software e.g. CheckM (Parks et al., 2015) or manually is strongly recommended.

## 3 Results

Metagenome binning performance of YAMB was tested on real metagenome of alkaline hot spring on desktop computer (CPU with 4 threads, 16GB RAM, Ubuntu 16.04). Metagenome was assembled using SPAdes (Bankevich, A. et al., 2012). Sequencing data from alkaline thermal spring (NCBI SRA: SRX3521399 and SRX3521400) was assembled to 24 578 contigs with total length 31 020 002 bp, 7 040 contigs had length of at least 1000 bp (total length 30 075 232 bp). Reads from each sequencing run were mapped using bowtie2 (Langmead, B., Salzberg, S. L., 2012) and processed individually. Up to 18 bins were formed by YAMB depending on mapped perplexity parameter of t-SNE. The best binning case was selected by minimum share of unbinned sequences. Bin completeness and contamination assessed using CheckM (Parks, 2015). Metagenomic bins with completeness more than 50% are shown in table 1. YAMB binnnig results are comparable to the results of binning using CONCOCT on the same data in terms of completeness and contamination of metagenomics bins (from 77.21 to 98.78% and from 0.92 to 3.39% respectively for five most complete bins) (Korzhenkov A. et al., 2018). Provisional taxonomical attribution was performed using NCBI BLAST (Johnson, M. et al., 2008) in blastx mode.

**Table 1.**
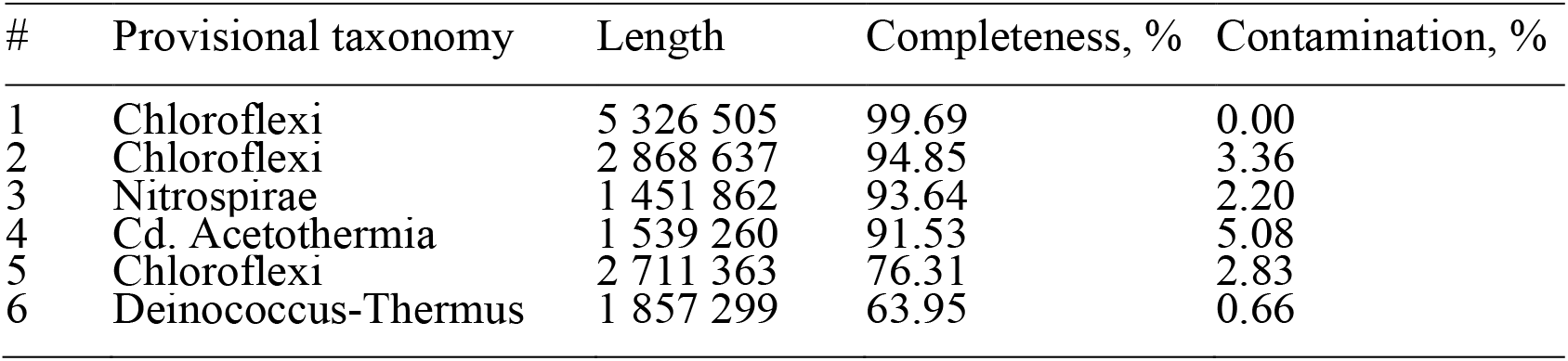
Metagenomic bins, discovered in the hot alkaline spring metagenome

## 4 Conclusion

YAMB shows itself as a successful application of modern methods of data dimensionality reduction and clustering – tSNE and HDBSCAN – to metagenome binning. YAMB may produce bins with high completeness and low contamination from real sequencing data.

## Acknowledgements

Author thanks Oleg Mozhey and Evgenii Lunev for results evaluation and useful comments.

